# An imaging flow cytometry method to study platelet-monocyte aggregates using Long COVID as a model

**DOI:** 10.64898/2026.04.09.717442

**Authors:** Anél Thompson, Chantelle Venter, Willem JS de Villiers, Dalene De Swardt, Gert J Laubscher, Douglas B Kell, Etheresia Pretorius

**Affiliations:** Department of Physiological Sciences, Faculty of Science, Stellenbosch University, Stellenbosch Private Bag X1 Matieland, 7602, South Africa; Department of Gastroenterology, Faculty of Medicine and Health Sciences; Central Analytical Facility, Flow Cytometry Unit, Stellenbosch University, Tygerberg Campus, Room 2096, Biomedical Research Institute, Francie Van Zijl Drive, Tygerberg, Cape Town 7505, South Africa; Suite 104, 1 Elsie du Toit street, Mediclinic Stellenbosch, Stellenbosch 7600, South Africa; Department of Biochemistry, Cell and Systems Biology, Institute of Systems, Molecular and Integrative Biology, Faculty of Health and Life Sciences, University of Liverpool, Crown St, Liverpool L69 7ZB, UK

**Keywords:** Long COVID, Thromboinflammation, Platelet-monocyte aggregates, Imaging Flow Cytometry

## Abstract

**Background:** Long COVID is characterised by persistent systemic inflammation and endothelial dysfunction, with increasing evidence implicating thromboinflammatory mechanisms. Platelet-monocyte aggregates (PMA) represent a sensitive marker of platelet activation and immune-vascular interactions, but their role in Long COVID remains incompletely defined.

**Methods:** This study quantified circulating PMA in 20 Long COVID patients and 20 healthy controls using a two-colour imaging flow cytometry assay targeting CD14 (a monocyte receptor for pathogen-associated molecular patterns, PAMPs) and CD62P (P-selectin). PMA were expressed as a percentage of total monocytes, and platelet attachment patterns were classified into single versus multiple platelet binding. Statistical analyses included Shapiro-Wilk normality testing, unpaired t-tests, Mann-Whitney U tests or two-way ANOVA as appropriate, and linear regression for correlation analysis.

**Results:** Circulating PMA were significantly elevated in Long COVID patients compared with controls (29.19 [20.02–37.26] vs 4.59 [2.67–7.16], p < 0.0001). Long COVID samples showed a reduced proportion of monocytes with single platelet attachment and a corresponding increase in multiple platelet binding (p < 0.0001). In controls, %PMA increased with age (p < 0.01), whereas no age association was observed in Long COVID, indicating an elevated baseline independent of age.

**Conclusions:** Long COVID is associated with markedly increased platelet–monocyte aggregation and altered platelet attachment dynamics, consistent with sustained thromboinflammatory activity. PMA represent a sensitive cellular marker of platelet-driven immune activation and may have utility as an accessible biomarker for stratifying thromboinflammatory burden in Long COVID.

## Introduction

During the early phase of the COVID-19 pandemic, it became evident that a subset of individuals infected with severe acute respiratory syndrome coronavirus 2 (SARS-CoV-2) experienced a broad spectrum of persistent symptoms that continued well beyond the resolution of the acute infection (1-7). This condition, known as Long COVID or post-acute sequelae of COVID-19 (PASC), has since been recognised as a complex, multisystemic syndrome associated with numerous clinical and public health challenges (8-13). Long COVID is estimated to affect 4% of children and 10–26% of adults following SARS-CoV-2 infection (11), and with over 651 million documented cases of COVID-19 worldwide, the global burden is estimated at a minimum of 65 million individuals (14).

Current evidence suggests that in Long COVID, persistent systemic inflammation and endothelial dysfunction reinforce one another through ongoing thromboinflammatory mechanisms (13, 15-20). One such mechanism involves continued immune activation following acute SARS-CoV-2 infection, resulting in a prolonged and dysregulated immune response (9, 17, 19, 21, 22). Elevated C-reactive protein (CRP) levels, consistent with chronic, low-grade inflammation, are associated with the presence of Long COVID symptoms (10) and promote oxidative stress via increased reactive oxygen species (ROS) production, thereby contributing to endothelial dysfunction (23).

Emerging evidence has identified fibrinaloid microclot complexes (FMCs) as a hallmark of thromboinflammatory pathology in Long COVID (12, 13, 17, 19, 20). These circulating, fibrinolysis-resistant structures are enriched in fibrin(ogen) and inflammatory molecules, as well as amyloidogenic proteins, and have been shown to persist in the plasma of affected individuals (12). FMCs are structurally associated with components of innate immune activation, including neutrophil extracellular traps, and are thought to contribute to microvascular dysfunction and impaired fibrinolysis (17). Within this framework, cellular interactions such as platelet-monocyte aggregate formation may represent an additional, complementary layer of thromboinflammatory activity.

Platelet-monocyte interactions have emerged as a key mechanistic interface linking thrombotic processes with inflammatory pathways (24), which may represent the cellular interface where immune activation and endothelial injury converge. Platelets are small, anucleate blood components that play crucial roles in haemostasis, as well as in immune processes that drive vascular inflammation (25). In Long COVID, platelets have been reported to exhibit a hyperactivated, aggregate prone phenotype (12, 26). Activated platelets can attach to circulating monocytes via platelet P-selectin and monocyte P-selectin glycoprotein ligand-1 (PSGL1) to form platelet-monocyte aggregates (PMA) (27). Circulating PMA levels have been found to be elevated in various thromboinflammatory and autoimmune conditions, including cardiovascular diseases (28-30), sepsis (31), acute COVID-19 (32-34), type 1 diabetes (35), systemic lupus erythematosus, and rheumatoid arthritis (36), highlighting them as promising biomarkers of immune-vascular dysfunction.

As persistent systemic inflammation drives increased formation of circulating PMA, reflecting thromboinflammatory processes across inflammatory diseases, the aim of this study was to quantify circulating PMA levels in Long COVID patients compared with healthy controls. These measurements may support the use of PMA as one of the accessible cellular biomarkers for stratifying the thromboinflammatory burden in Long COVID.

## Methods and materials

### Ethical clearance

This study obtained ethical clearance from the Health Research Ethics Committee (HREC) of Stellenbosch University (South Africa) (HREC Reference No: S24/07/181, Project ID: 31404). All samples were obtained via the Stellenbosch University Biorepository (Project ID: 28804) and were fully anonymized prior to use. Prior to blood sample collection, all volunteers were fully informed about the study’s objectives, potential risks, procedural details and informed consent was obtained from each study participant. The study adhered strictly to ethical guidelines, including the Declaration of Helsinki, the South African Guidelines for Good Clinical Practice and the Medical Research Council Ethical Guidelines for Research.

### Blood sample collection

Fresh whole blood was collected via venepuncture into 2.7 mL sodium citrate (3.2%) vacuum tubes (BD Vacutainer®, 363048), ensuring a 1:9 citrate-to-blood volume ratio. Each tube was gently inverted immediately after collection to prevent clotting and ensure thorough mixing. All blood collections adhered to standard sterile protocol and were performed by a certified phlebotomist or medical professional.

### Patient demographics

The study included 20 healthy individuals to serve as controls (9 males; 11 females; median age: 25 [22–35]) and 20 Long COVID patients (7 males; 13 females; median age: 45 [37–52]). Study participants were not recruited based on age- or sex-matching criteria. All participants completed a comprehensive questionnaire that gathered information regarding general demographics and clinical history such as pre-existing co-morbidities along with their self-reported Long COVID symptoms. Healthy controls were defined as individuals with no known history of coagulopathies, were not pregnant, and had no acute infections or symptoms associated with Long COVID at the time of recruitment. All controls were non-smokers, with the except of one individual who was otherwise clinically healthy with no comorbidities. One control participant reported gut dysbiosis and elevated cholesterol levels. Long COVID participants were identified based on self-reported persistent symptoms consistent with post-acute sequelae of SARS-CoV-2 infection. While all 20 participants included in this study exhibited symptom profiles consistent with Long COVID and were referred to us by our clinical collaborator, who diagnosed them with Long COVID according to the WHO criteria, 11 reported a confirmed prior COVID-19 diagnosis, while 7 reported no testing history and 2 reported no confirmed diagnosis, but suspect in infection. All participants reported that their symptoms arose after a confirmed or suspected acute infection and symptoms lasted for at least 3 months and were not present prior to infection or suspected infection. **Table 1** provides an overview of the median age and gender distribution of the control and Long COVID groups, as well as a detailed summary of the Long COVID participants’ co-morbidities and symptoms.

**Table 1.** Sample Demographics.

| **Median age [and interquartile range (Q1–Q3)] of controls and Long COVID patients** |  |
| --- | --- |
| Controls (n=20) | 25 [22–35] |
| Long COVID (n=20) | 45 [37–52] |
| **Gender of controls and Long COVID patients** |  |
| Controls (n=20) | 9 males; 11 females |
| Long COVID (n=20) | 7 males; 13 females |
| **Vaccination status of controls and Long COVID patients before blood collection** |  |
| Percentage of Controls vaccinated (n=20) | 65% |
| Percentage of Long COVID vaccinated (n=20) | 75% |
| **Co-morbidities of Long COVID patients (n=20)** |  |
| **Co-morbidity** | **% In 20 Long COVID patients** |
| Gut dysbiosis | 15% |
| High cholesterol | 15% |
| ME-CFS | 15% |
| Periodontitis or Gingivitis | 10% |
| High blood pressure | 10% |
| Cancer | 10% |
| Psoriasis | 10% |
| Rosacea | 5% |
| Eczema | 5% |
| Fibromyalgia | 5% |
| Hypothyroidism | 5% |
| **Symptoms of Long COVID patients (n=20)** |  |

| Symptom | % In 20 Long COVID patients |
| --- | --- |
| Constant fatigue | 95% |
| Brain fog/Concentration issues | 85% |
| Depression/Anxiety | 75% |
| Shortness of breath | 75% |
| Sleep disturbances | 65% |
| Post exertional malaise | 60% |
| Heart palpitations | 60% |
| Digestive issues | 60% |
| Tinnitus | 50% |
| Joint and muscle pains | 50% |
| Chest pains | 35% |
| Headaches | 20% |

### Panel design

The panel design consists of a two-colour platelet-monocyte aggregate assay (**Table 2**), which forms the basis for analysis by imaging flow cytometry. Included in the panel is a Brilliant Violet 421-(BV421-) conjugated mouse anti-human CD14 monoclonal antibody (clone 61D3, 404-0149-42, eBioscience™) and a Phycoerythrin- (PE-) conjugated mouse anti-human CD62P monoclonal antibody (clone CLBThromb/6, IM1759U, Beckman Coulter Life Sciences). CD14 serves as a marker for the lipopolysaccharide (LPS) receptor, which is highly expressed on the surface of monocytes (37). CD62P is a marker for P-selectin, which is found within platelet α-granules (38) or located on platelet surface membranes following translocation from granules as a result of platelet activation (39). Acquisition settings were optimised for each fluorophore based on their distinct spectral excitation and emission profiles (**Table 2**) ensuring accurate detection of each marker.

**Table 2.** Two-colour platelet-monocyte aggregate assay.

|  | CD14 BV421 | CD62P PE |
| --- | --- | --- |
| **Excitation $\lambda$** | 406nm | 488nm |
| **Emission $\lambda$** | 420-505 nm | 505-560 nm |

### Sample preparation

Sample preparation involved staining 200 µL of whole blood with 10 µL CD14 BV421 (final concentration of 4.65 µg/mL) and 5 µL CD62P PE (final concentration of 0.145 µg/mL). Samples were incubated for 30 minutes at room temperature in the dark. Fixation and red blood cell lysis was performed with 2 mL of 10× diluted BD FACS™ Lysing Solution (BD Biosciences, USA), and the samples were incubated for an additional 15 minutes at room temperature.

After fixation and lysis, samples were centrifuged at 200×g for 5 minutes and the supernatant was removed. The remaining volumes were washed with 1 mL of phosphate-buffered saline (PBS, Gibco™) and centrifuged again under the same conditions. After discarding the supernatant, the samples were resuspended in 100 µL of PBS for imaging flow cytometry analysis.

### Sample acquisition and analysis

A FlowSight® Imaging Flow Cytometer from Cytek® Biosciences was used to analyse samples with the INSPIRE v200.0.336.0 acquisition software (Amnis®). The excitation lasers used included the 100 mW 405 nm and 60 mW 488 nm lasers. CD14 BV421 was excited by the 405 nm laser, with emission detected between 420 and 505 nm (Channel 07), while CD62P PE was excited by the 488 nm laser, with emission detected between 505 and 560 nm (Channel 03) (**Fig. 1**). A 12 mW 785 nm laser was used for side scatter measurements (Channel 06). All samples were acquired at a low flow rate, and images were captured with a 20 × objective in the brightfield channel (Channel 01).

**Figure 1.**
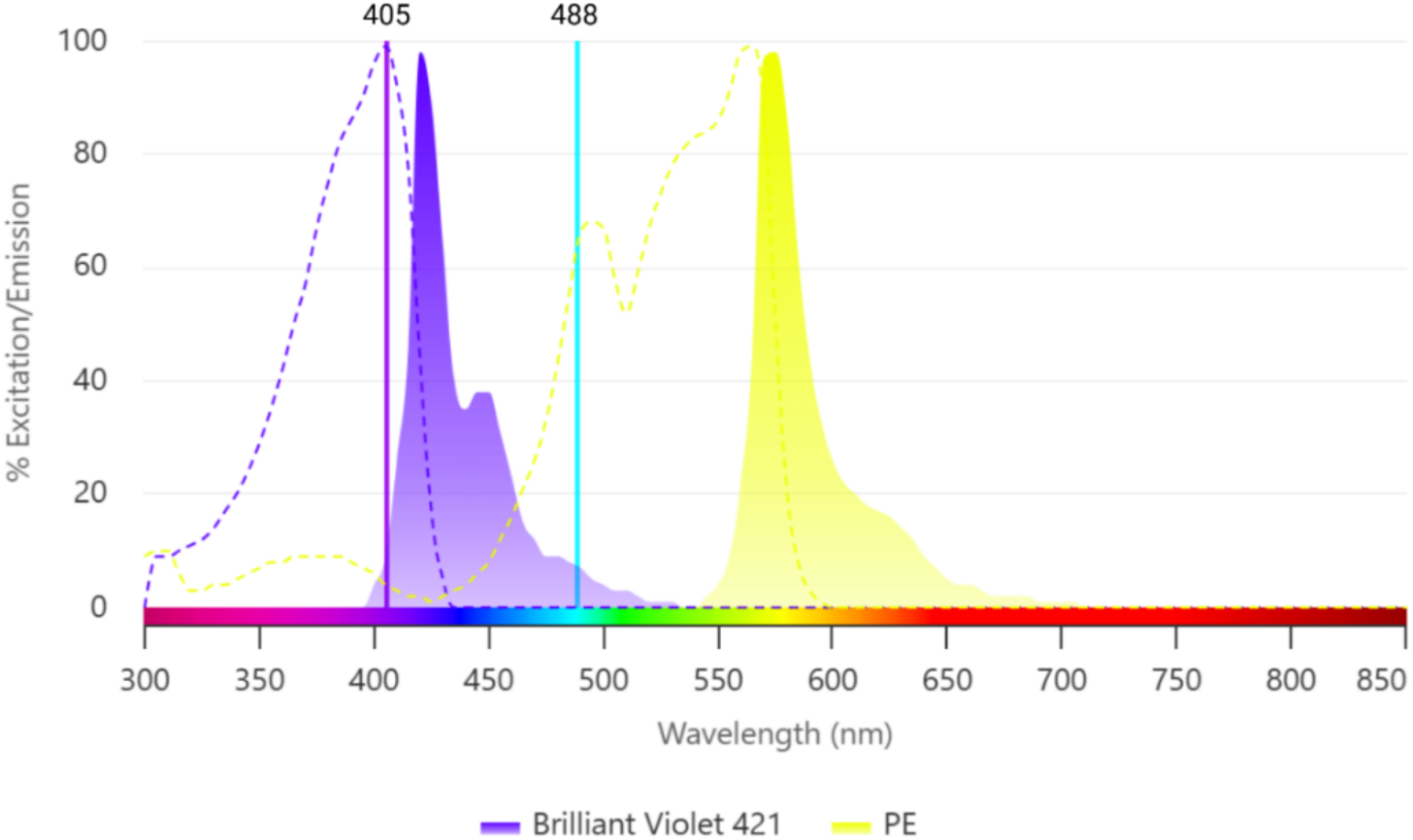
Spectral map of platelet-monocyte aggregate assay. The combined fluorophores were excited by a dual laser setup with two different laser wavelengths: the 405nm laser for CD14 BV421 and the 488nm laser for CD62P PE. The dotted lines represent the excitation spectra, while the solid lines represent the emission spectra for each antibody. This figure was generated using the Fluorescence Spectrum Analyzer for Flow Cytometry by Beckman Coulter Life Sciences (https://www.beckman.co.za/flow-cytometry/fluorescence-spectrum-analyzer).

Imaging flow data analysis was performed with the IDEAS® v6.4 image analysis software (Amnis®). A total of 2000 monocytes, identified by their characteristic side scatter properties and positive staining for CD14 (Ch07) (**Fig. 2A**), were counted per sample with platelet-monocyte aggregates reported as a percentage of the counted monocytes. Cell doublets, debris, and aggregates were excluded from the analysis by applying a brightfield area versus aspect ratio gate around single monocytes (with or without attached platelets), which have a high aspect ratio and intermediate area (**Fig. 2B**). CD62P fluorescence intensity (Ch03) was used to distinguish CD62P negative from CD62P positive events, and a gate was drawn around the CD62P^+^ population, representing monocytes with attached platelets (**Fig. 2C**).

**Figure 2.**
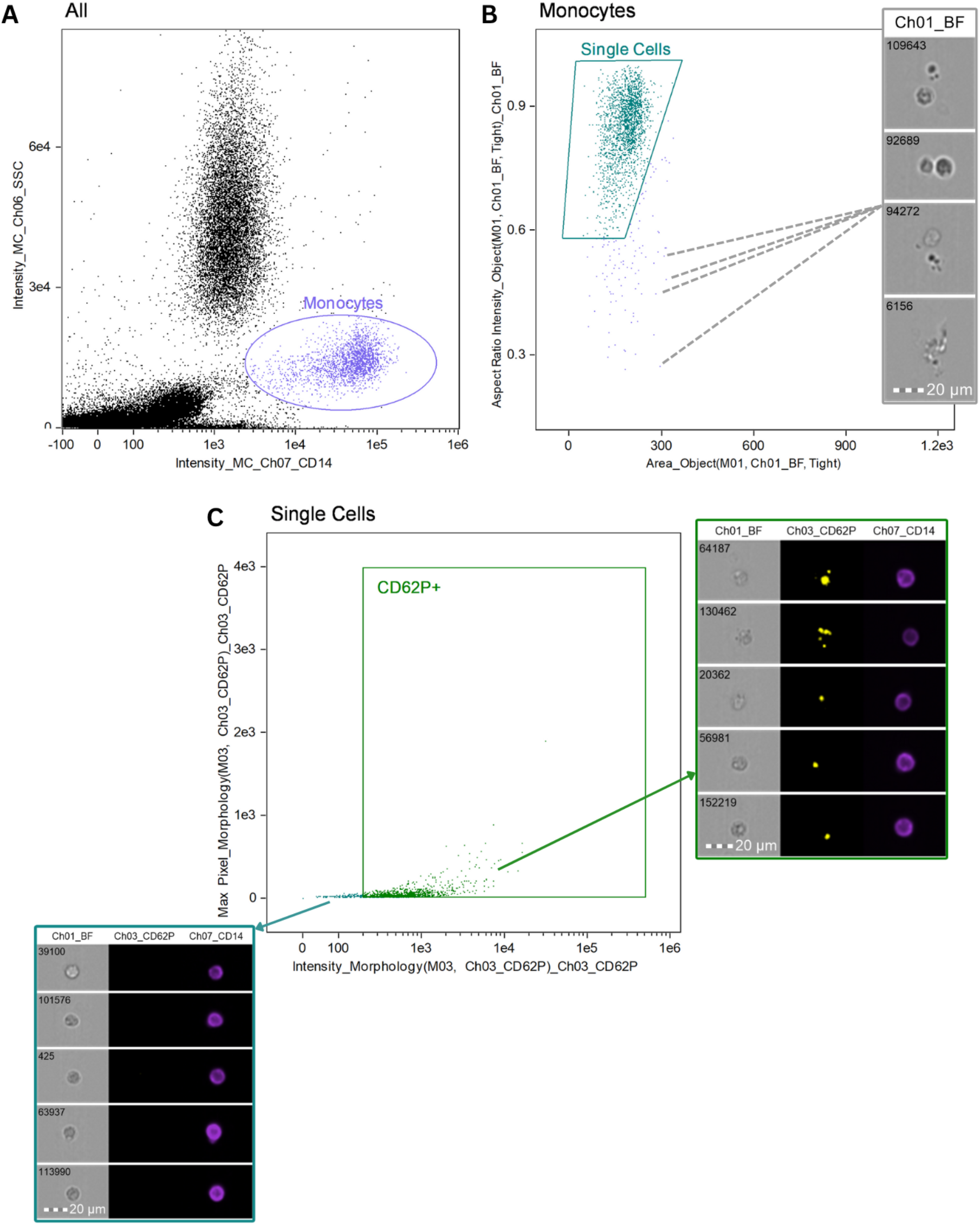
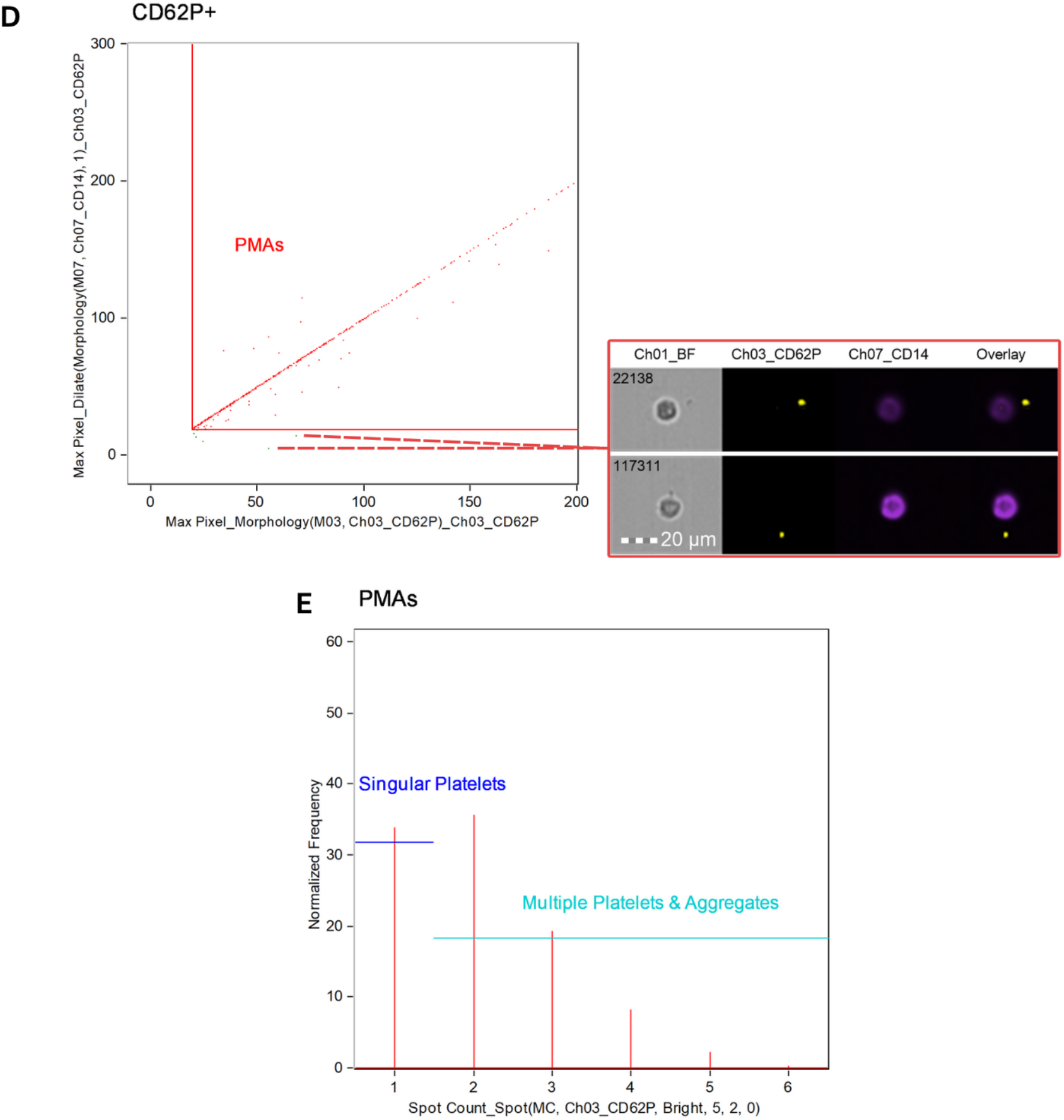
Gating strategy for identifying platelet-monocyte aggregates using imaging flow cytometry. **(A)** Monocytes were identified based on CD14 fluorescence intensity (Ch07) and intermediate side scatter properties. **(B)** Single cells were gated using brightfield (Ch01) area versus aspect ratio to exclude debris, aggregates, and cell doublets (grey box). **(C)** Within single monocytes, CD62P fluorescence intensity (Ch03) was used to distinguish platelet positive events (green box) from monocytes without attached platelets (blue box). Platelet–monocyte aggregates were defined as CD14^+^CD62P^+^ events, with the red gate used to distinguish true aggregates from coincidence events (red box). **(E)** Spot count analysis of CD62P fluorescence was used to quantify the number of platelets per monocyte, distinguishing monocytes with a single platelet from those with multiple or aggregated platelets. Channel 1 shows brightfield; Channel 3 shows CD62P fluorescence; Channel 7 shows CD14 fluorescence; and the overlay combines the Ch03 and Ch07 signals.

Coincidence events, events that occur when platelets and monocytes pass through the flow detector simultaneously but are not truly associated, were excluded from the CD14^+^CD62P^+^ population. A dilated (plus 1 pixel) CD14 morphology mask (**Fig. S1**) was used to define the monocyte signal and the area immediately around the cell. The max pixel intensity of CD62P was then evaluated exclusively within the dilated CD14 mask, restricting CD62P signal assessment to regions corresponding to the monocyte surface area. CD62P fluorescence detected outside the mask, which reflects non-attached platelets and coincidence events, was excluded **(Fig. 2D)**. To determine the number of platelets bound per monocyte, a spot count mask was applied to the CD62P signal, generating individual spots for each platelet within a single monocyte event. Gates were created to distinguish monocytes with a single platelet attached from those with multiple platelets or platelet aggregates **(Fig.2E)**. Representative images of monocytes with single or multiple/aggregated platelets attached are shown in **Supplementary Figure S2**.

### Confocal microscopy

Sample preparation followed the same protocol as described for imaging flow cytometry, that was followed by pipetting 10uL of the sample onto a glass slide and covered with a coverslip for imaging. Samples were viewed with the Evident Microscopy Fluoview FV4000 Confocal Laser Scanning Microscope (Tokyo, Japan) using the UPLXAPO 60x objective. To visualize the CD14 BV421 (purple fluorescence), the 405nm laser was used with the emission wavelength set at 430-470nm. For the CD62P PE (yellow fluorescence) the 561nm laser was used with the excitation wavelength set at 570-620nm.

### Statistical analysis

Statistical analyses were performed using GraphPad Prism 10 (version 10.6.1), with Microsoft® Excel® used for data organisation. Data normality was assessed using the Shapiro–Wilk test. For normally distributed data, differences between two groups were evaluated using an unpaired t-test, while comparisons involving two independent variables were analysed using two-way ANOVA with Šidák-corrected multiple comparisons where applicable. Normally distributed data are presented as mean ± standard deviation. For non-normally distributed data, the Mann–Whitney U test was used, with results presented as median [Q1–Q3]. All statistical tests were two-tailed. Correlation analysis was performed using simple linear regression. Categorical variables were compared using Fisher’s exact test. Statistical significance was defined as p < 0.05.

## Results

### Participant features

Our study included a total of 40 participants, comprising 20 healthy controls and 20 Long COVID patients. Participant demographics are summarised in **Table 1**. An unpaired, two-tailed Mann–Whitney test showed that participants in the Long COVID group (median age 45 years [IQR 37–52]) were significantly older than those in the control group (median age 25 years [IQR 22–35]; p < 0.01). The control group comprised 55% females and 45% males, while the Long COVID group included 65% females and 35% males. A two-sided Fisher’s exact test indicated that the gender distribution did not differ significantly between groups (p ≥ 0.05). A two-way ANOVA assessed the effects of group (control vs Long COVID), sex (female vs male), and their interaction, with Šidák-corrected multiple comparisons used for within-sex and within-group comparisons. Long COVID participants exhibited significantly higher %PMA compared with controls in both females (p < 0.0001) and males (p < 0.001), while no significant sex differences in %PMA were observed within either group (p ≥ 0.05) (**Fig.3**).

**Figure 3.**
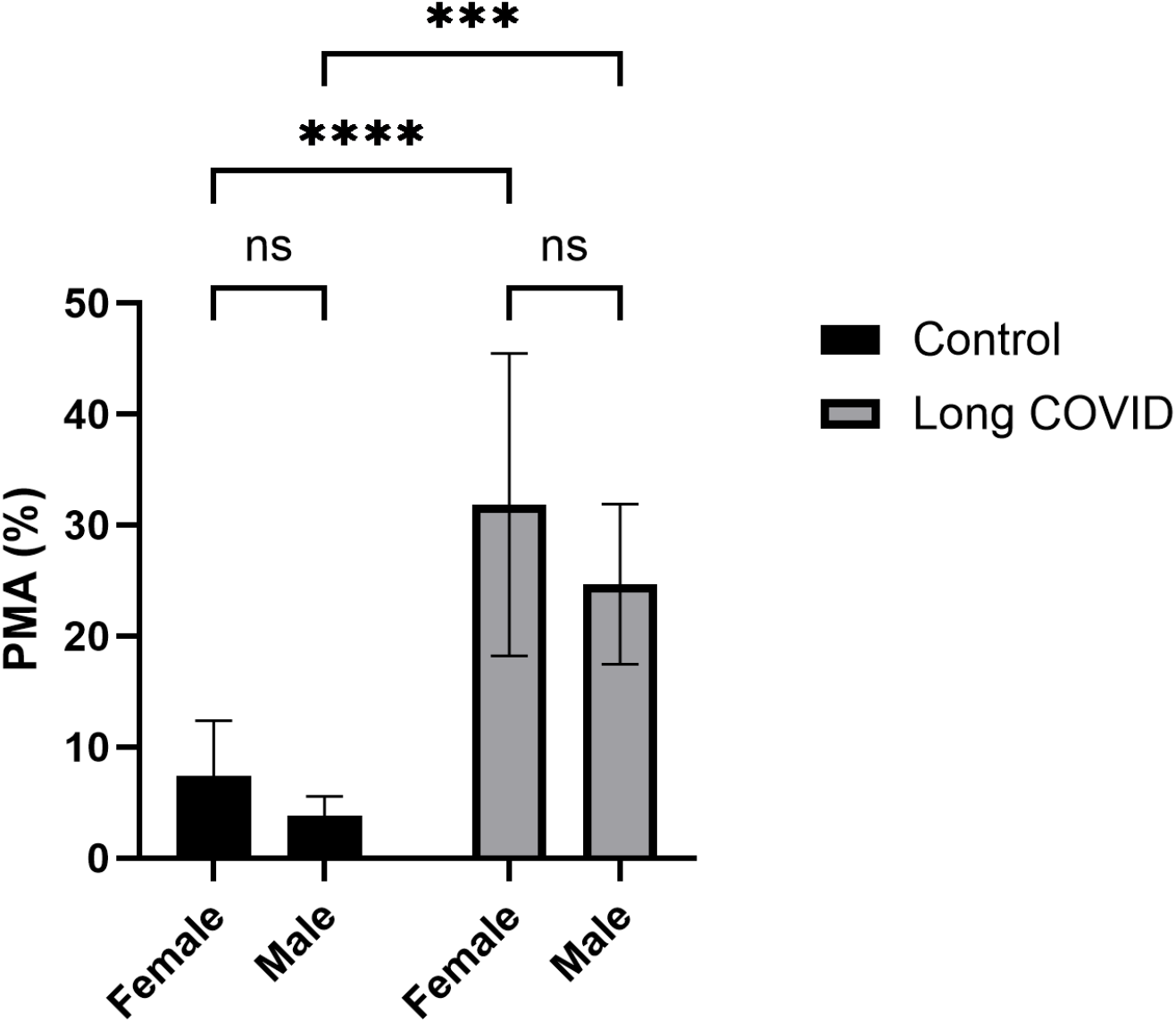
Percentage PMA in control and Long COVID participants stratified by sex. Data are presented as mean ± SD. A Two-way ANOVA with Šidák-corrected multiple comparisons showed no significant sex-based differences in %PMA within either the control or Long COVID groups. ns, not significant; ***p<0.001; ****p<0.0001.

The Long COVID participants reported a range of pre-existing co-morbidities, with gut dysbiosis, high cholesterol, and myalgic encephalomyelitis/chronic fatigue syndrome (ME-CFS) each reported by 15% of patients, and periodontal disease, hypertension, cancer, and psoriasis reported by 10% of patients. Symptom burden was high in the Long COVID group, with persistent fatigue reported by 95% of participants. Additionally, more than 70% of patients also reported cognitive impairment (“brain fog”), depression or anxiety, and shortness of breath.

Simple linear regression (**Fig.4**) showed that %PMA in healthy controls increased somewhat with age (slope = 0.178, p < 0.01, R^2^ = 0.32). Importantly, however, the %PMA in Long COVID samples showed no significant relationship with age (slope = 0.224, p ≥ 0.05, R^2^ = 0.05). Comparison of control and Long COVID regression lines showed that the slopes were not significantly different (p ≥ 0.05), indicating a similar rate of PMA increase with age, whereas the intercepts were significantly higher in Long COVID samples (p < 0.0001), reflecting an elevated baseline %PMA independent of age. Thus, the increased levels of PMA in Long COVID patients that we observe here is not an artefact related to age.

**Figure 4.**
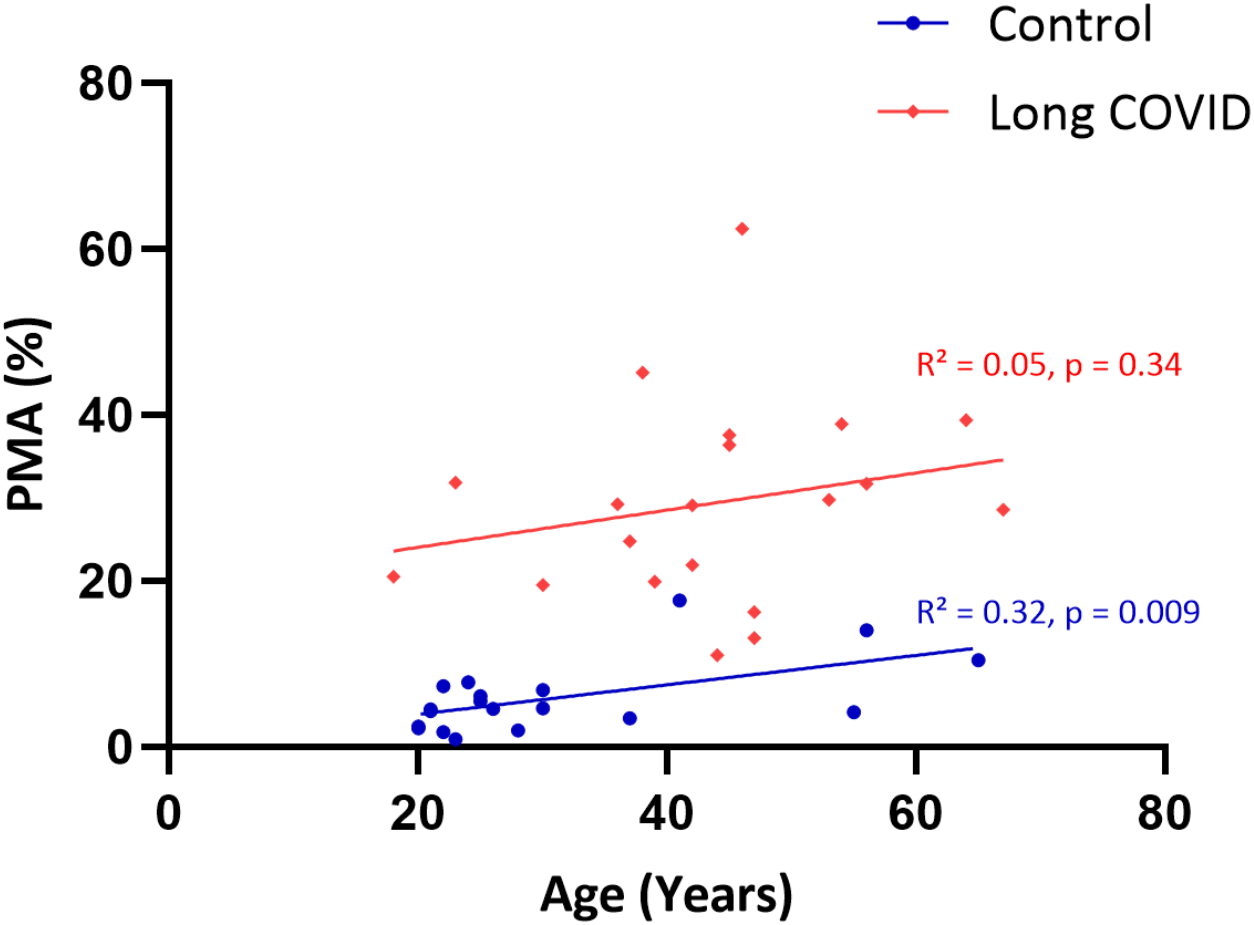
Scatter plot showing the relationship between %PMA and age in control and Long COVID participants. Each point represents an individual participant. Solid blue circles and regression line indicate the control group, while red diamonds and regression line represent the Long COVID group. Simple linear regression revealed a significant positive correlation between age and %PMA in controls (slope = 0.178, R^2^ = 0.32, p < 0.01), whereas no significant correlation was observed in Long COVID samples (slope = 0.224, R^2^ = 0.05, p ≥ 0.05, ns). PMA, Platelet-monocyte aggregates.

### Elevated %PMA and altered platelet attachment patterns in Long COVID

Representative images of PMA from both control and Long COVID samples are shown in **Figure 5**. A two-tailed unpaired Mann–Whitney test showed that circulating %PMA were significantly elevated in the Long COVID group compared with controls (29.19 [20.02–37.26] vs. 4.59 [2.67–7.16], p < 0.0001) (**Fig.6A**). A two-tailed unpaired Welch’s t-test showed that the proportion of monocytes with a singular platelet attached was significantly lower in Long COVID samples than in controls (41.99 [±15.75] vs. 69.93 [±11.80], p < 0.0001) (**Fig.6B**). Similarly, a two-tailed unpaired Welch’s t-test showed that the proportion of monocytes with multiple/aggregated attached platelets was significantly higher in Long COVID samples than in controls (57.41 [±15.80] vs. 29.79 [±11.81), p < 0.0001) (**Fig.6C**).

**Figure 5.**
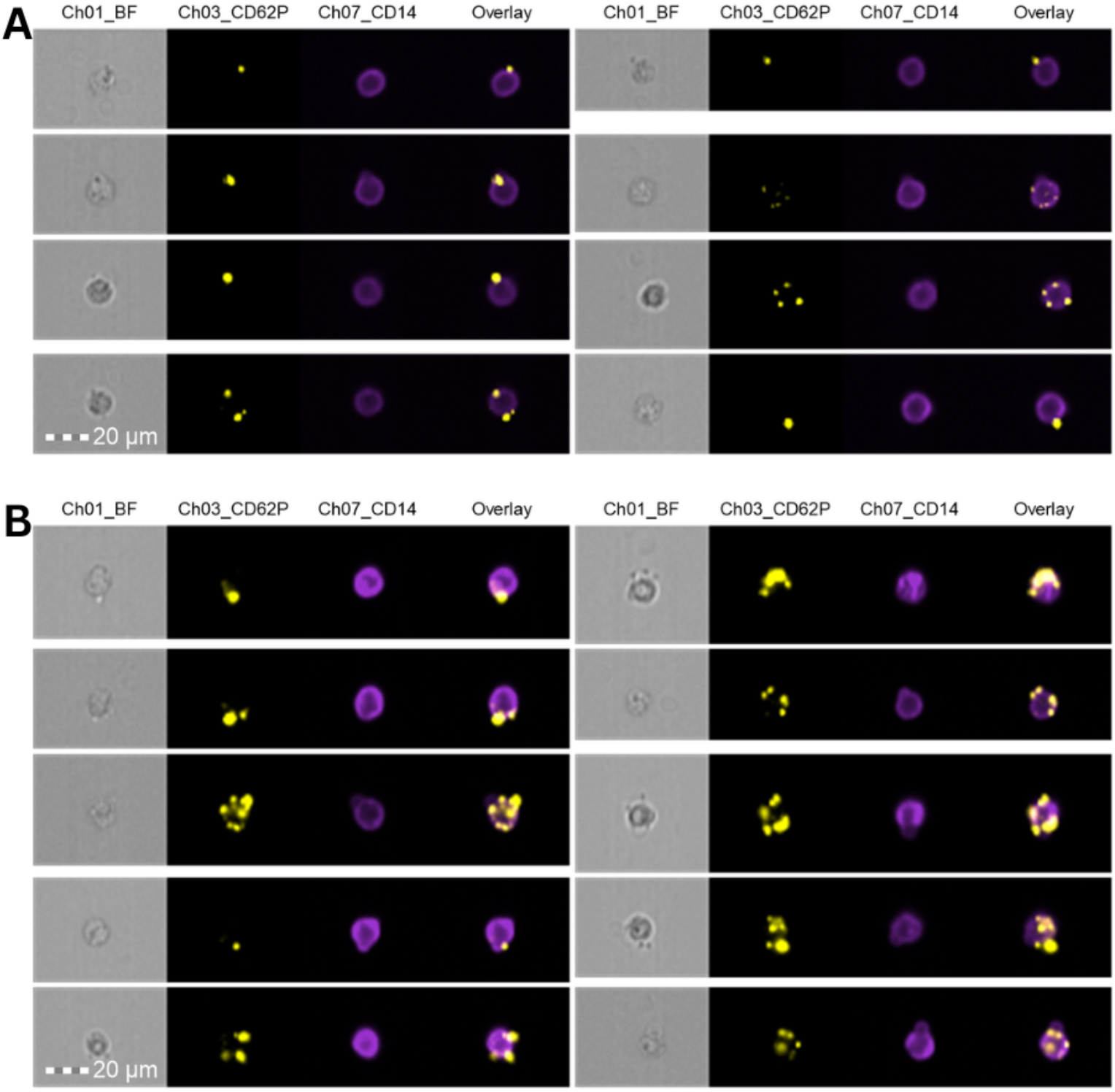
Representative images of **(A)** control and **(B)** Long COVID PMA. Channel 1 shows brightfield; Channel 3 shows CD62P fluorescence; Channel 7 shows CD14 fluorescence; and the overlay combines the Ch03 and Ch07 signals.

**Figure 6.**
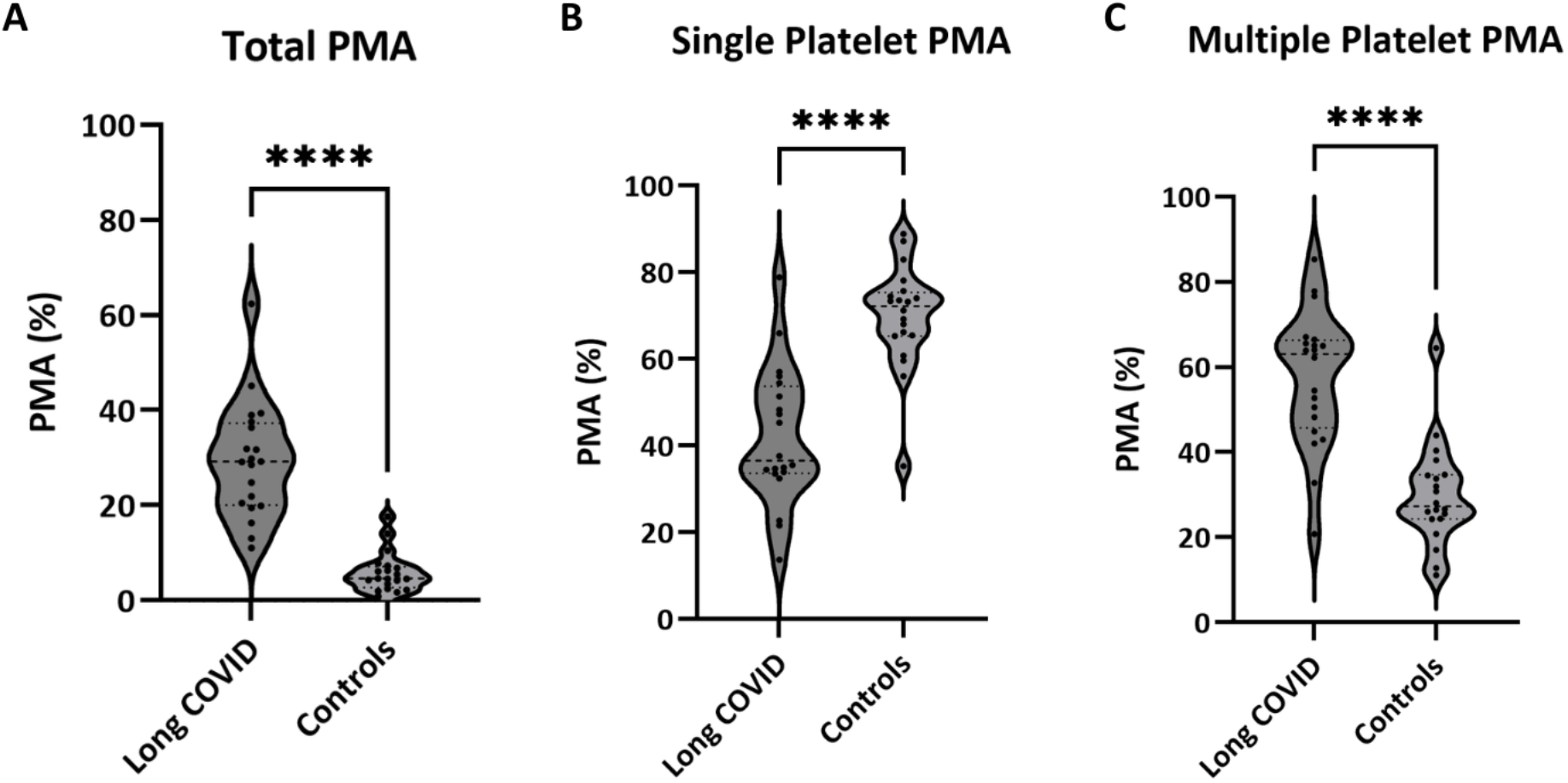
%PMA and platelet attachment phenotypes in Long COVID and control participants. (**A**) Total %PMA per participant. (**B**) Proportion of monocytes with a single attached platelet. (**C**) Proportion of monocytes with multiple attached platelets. Box-and-violin plots show data distribution (violin), with median indicated by a dashed line and interquartile range (IQR) indicated by dotted lines. Statistical comparisons were performed using a two-tailed unpaired Mann–Whitney test for total %PMA and two-tailed unpaired Welch’s t-tests for singular and multiple platelet aggregates. **** indicates p < 0.0001.

### Confocal microscopy

Figure 7 shows representative confocal microscopy images demonstrating PMA in control and Long COVID samples, with qualitative differences in platelet attachment patterns. These images qualitatively demonstrate platelet attachment to CD14^+^ monocytes, including both single platelet binding and multiple platelet interactions.

**Figure 7.**
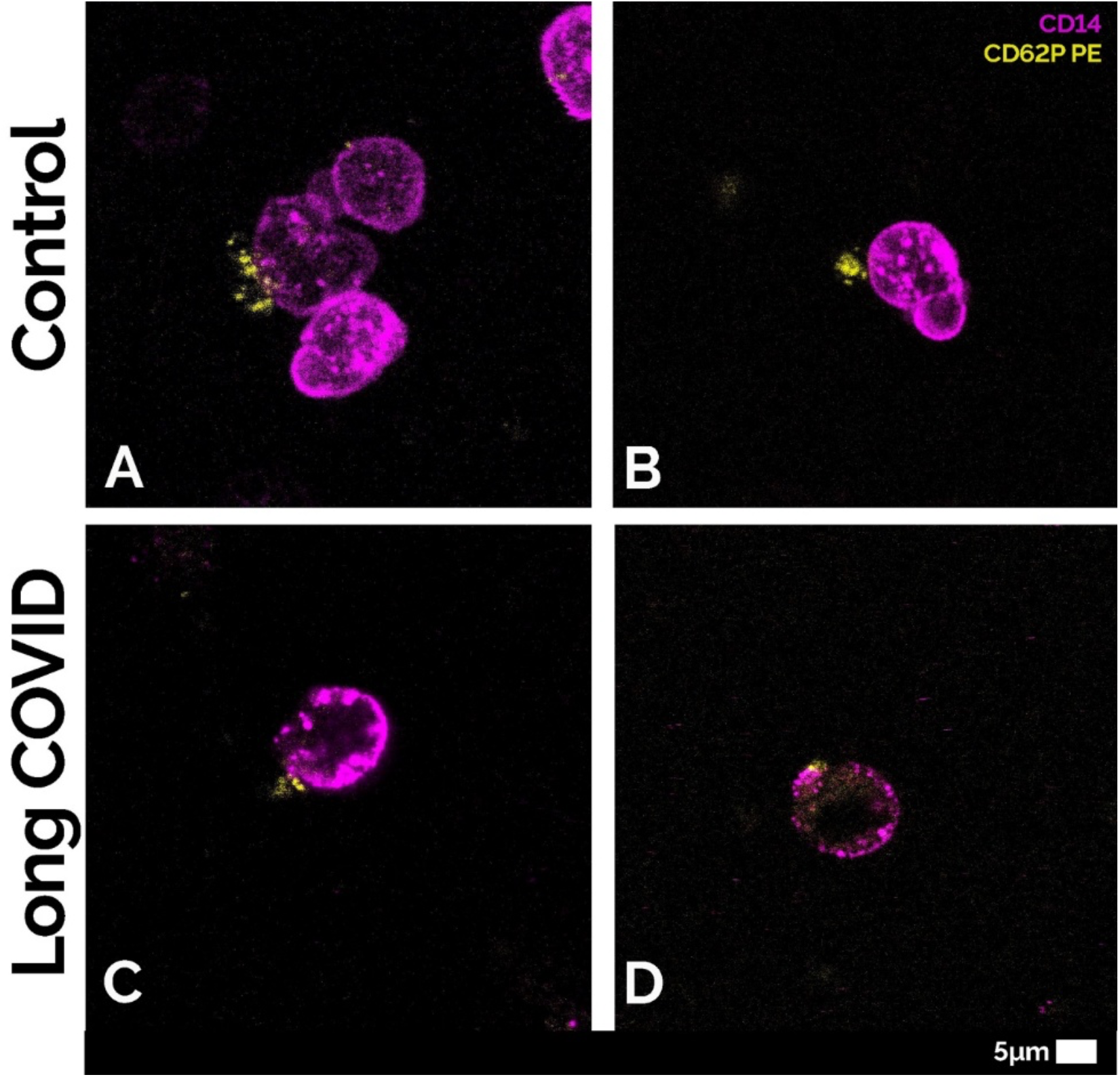
Representative images of control (**A and B**) and Long COVID (**C and D**) platelet–monocyte aggregates (PMA), that highlight the interactions between platelets and monocytes.

## Discussion

This study demonstrates a marked increase in circulating PMA in individuals with Long COVID, accompanied by a shift in platelet attachment patterns toward increased multiple platelet binding per monocyte. The findings of this study support the presence of sustained thromboinflammatory activity as well as persistent platelet activation in Long COVID and position PMA as a sensitive cellular marker reflecting this process. These findings also align with those reported by Brambilla et al.(10), who observed that Long COVID patients, relative to asymptomatic COVID-recovered subjects, display a platelet activation phenotype marked by elevated circulating platelet–leukocyte aggregates. These aggregates were further linked to residual lung abnormalities and the persistence of symptoms such as dyspnea, chest pain, and fatigue.

Circulating PMA are increasingly recognised as a highly sensitive indicator of platelet activation, often exceeding the sensitivity of platelet surface P-selectin expression alone (40). In healthy individuals, low levels of PMA are typically observed and may increase modestly with age (24, 41), reflecting baseline immune surveillance and vascular homeostasis. In our control population, %PMA increased with age and fell within the normal range of approximately 5%–20% reported for healthy individuals (24), and was consistent with values observed in control groups in other similar studies (42, 43). In contrast, Long COVID participants showed no significant association between age and %PMA, but rather a uniformly elevated baseline. This indicates that the increased PMA burden in Long COVID is not simply attributable to age but reflects a disease-associated shift in platelet–immune interactions.

Mechanistically, platelet–monocyte interactions represent a key interface between coagulation and immune activation (24, 42). Binding of activated platelets to monocytes via P-selectin– PSGL1 interactions promotes monocyte activation (44), upregulation of pro-inflammatory surface markers (45), and increased expression of tissue factor (34, 46), thereby enhancing thrombin generation and fibrin formation (47). In parallel, these interactions drive cytokine production and promote a pro-inflammatory monocyte phenotype, reinforcing a feed-forward loop between inflammation and coagulation (44, 45). The marked increase in PMA observed in this study therefore reflects not only platelet hyperactivation but also coordinated immune– vascular dysregulation.

Importantly, the shift from predominantly single platelet attachment in controls to increased multiple platelet binding in Long COVID suggests qualitative changes in platelet–monocyte interactions. Multiple platelet attachment may reflect enhanced platelet activation, increased platelet–platelet aggregation prior to monocyte binding or altered adhesive dynamics under inflammatory conditions. This phenotype is consistent with a more advanced or sustained thromboinflammatory state and may have functional consequences for monocyte activation and downstream coagulation pathways. Experimental models have shown that platelet binding can polarise monocytes toward a pro-inflammatory phenotype and enhance migratory and procoagulant behaviour (48).

These findings align with previous reports of platelet hyperactivation and thromboinflammatory pathology in Long COVID, including persistent endothelial dysfunction, increased inflammatory biomarkers, and platelet abnormalities (10, 12, 13, 26). Within this broader framework, PMA represent a complementary cellular readout of ongoing platelet-driven immune activation. Given their sensitivity and the relative simplicity of detection using flow-based platforms (37, 43), PMA may have potential as a translational biomarker for stratifying thromboinflammatory burden in Long COVID and related inflammatory conditions. The use of imaging flow cytometry in this study enables visualisation and morphological confirmation of PMA, thereby improving specificity by distinguishing true aggregates from coincident events.

These findings can be considered within the broader framework of FMCs, which have been identified as persistent, fibrinolysis-resistant structures in the circulation of individuals with Long COVID (12, 13, 17, 19). FMCs are enriched in inflammatory molecules and are structurally associated with components of innate immune activation, including neutrophil extracellular traps, suggesting a central role in sustaining thromboinflammatory processes (6, 7, 17). Within this context, elevated platelet-monocyte aggregates may represent a complementary cellular manifestation of the same underlying pathology. Platelet activation and aggregation contribute to the formation and stabilisation of fibrin(ogen)-rich microenvironments, while monocyte activation promotes tissue factor expression and amplifies coagulation cascades. The coexistence of increased PMA and FMCs therefore supports a model of persistent, self-reinforcing thromboinflammation in Long COVID, where cellular and molecular drivers of coagulation and inflammation are tightly coupled.

This study has several limitations. The sample size is modest, and the groups were not age-matched, although regression analysis supports an age-independent elevation of PMA in Long COVID. In addition, the cross-sectional design does not allow assessment of temporal dynamics or causal relationships. Future studies should include larger, longitudinal cohorts and integrate PMA measurements with molecular and proteomic profiling approaches to better define their relationship to disease severity, symptom clusters, and therapeutic response.

## Conclusion

In conclusion, Long COVID is associated with a substantial increase in circulating platelet– monocyte aggregates and a shift toward more complex platelet attachment patterns, consistent with sustained thromboinflammatory activity. These findings highlight PMA as a sensitive cellular marker of platelet-driven immune activation and support their potential utility as a biomarker for thromboinflammation in Long COVID and other inflammatory diseases (10, 28, 29, 31-33, 35, 36).

## Supporting information

All supplementary figures

## Author contributions

AT, EP, and WJSDV contributed to the conceptualisation of the study. AT, CV and DDS contributed to method development and data analysis. DBK, GJL and WJSDV reviewed and edited the manuscript.

## Consent for publication

Written informed consent was obtained from all the participants prior to the inclusion in the study.

