## Supplementary material for "An imaging flow cytometry method to study platelet-monocyte aggregates using Long COVID as a model": All supplementary figures

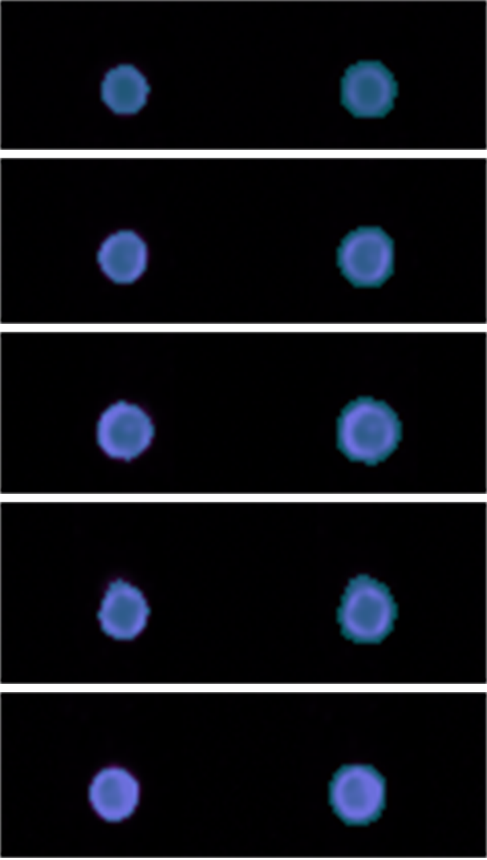

**Figure S1.** Dilated monocyte mask

*The figure illustrates the masking strategy applied to monocytes to exclude coincidence events. The left column shows the morphology mask overlaid on monocytes, used to identify monocytes based on fluorescence intensity. This mask fits tightly around the monocyte border, restricting the region of interest to the defined cellular boundaries. The right column shows the dilated (+1 pixel) mask overlaid on monocytes, which expands the monocyte region by one pixel to include the immediate space around the cell. This adjustment allows for the detection of CD62P signal from platelets attached to the monocyte surface.*

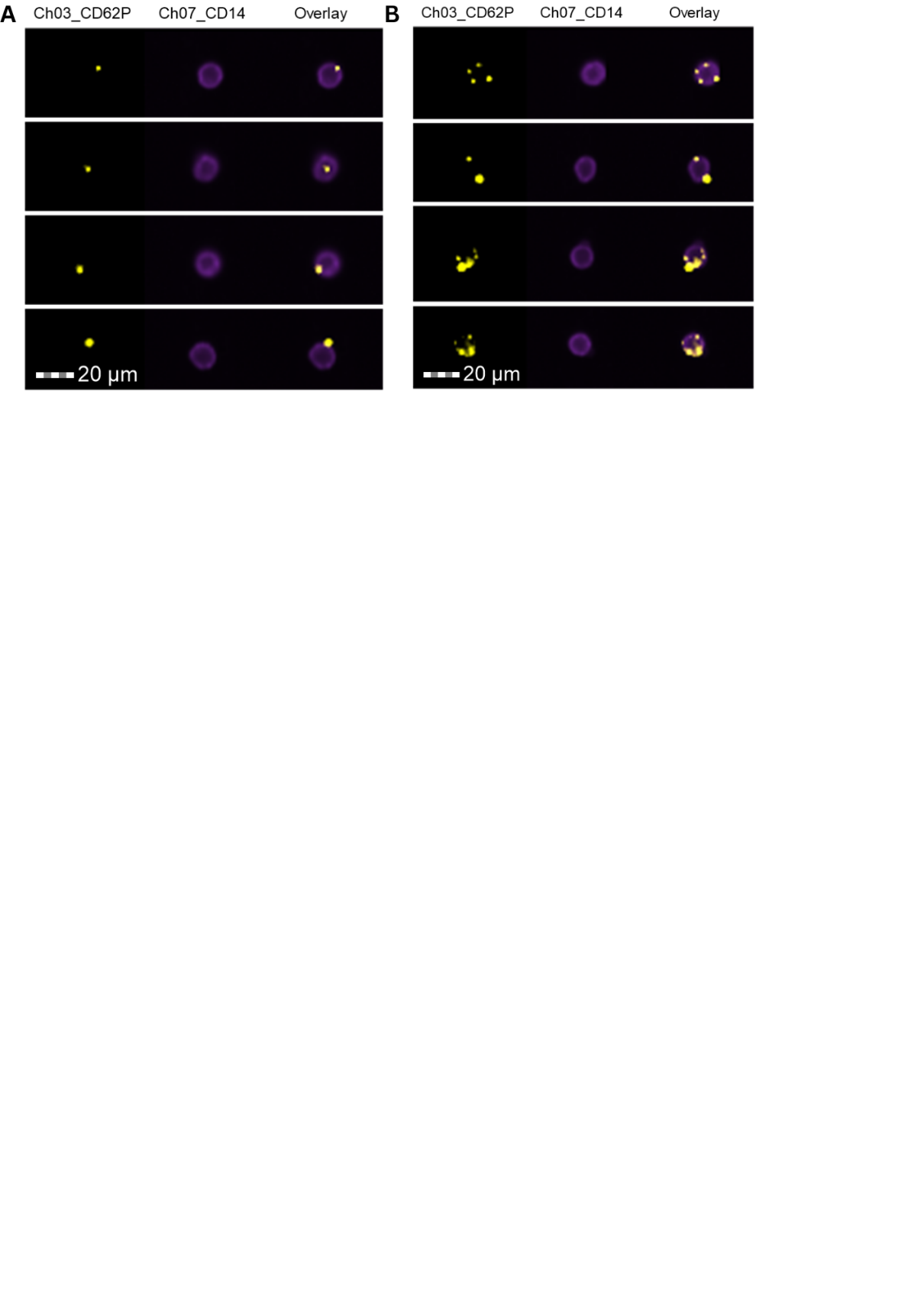

**Figure S2.** Representative images of monocytes with attached platelets

Representative images show **(A)** single platelets attached to monocytes and **(B)** multiple/aggregated platelets attached to monocytes, identified using the spot-counting feature. Channel 3 shows CD62P fluorescence; Channel 7 shows CD14 fluorescence; and the overlay combines the Ch03 and Ch07 signals.

**Appendix A: Ethical Clearance Letter**

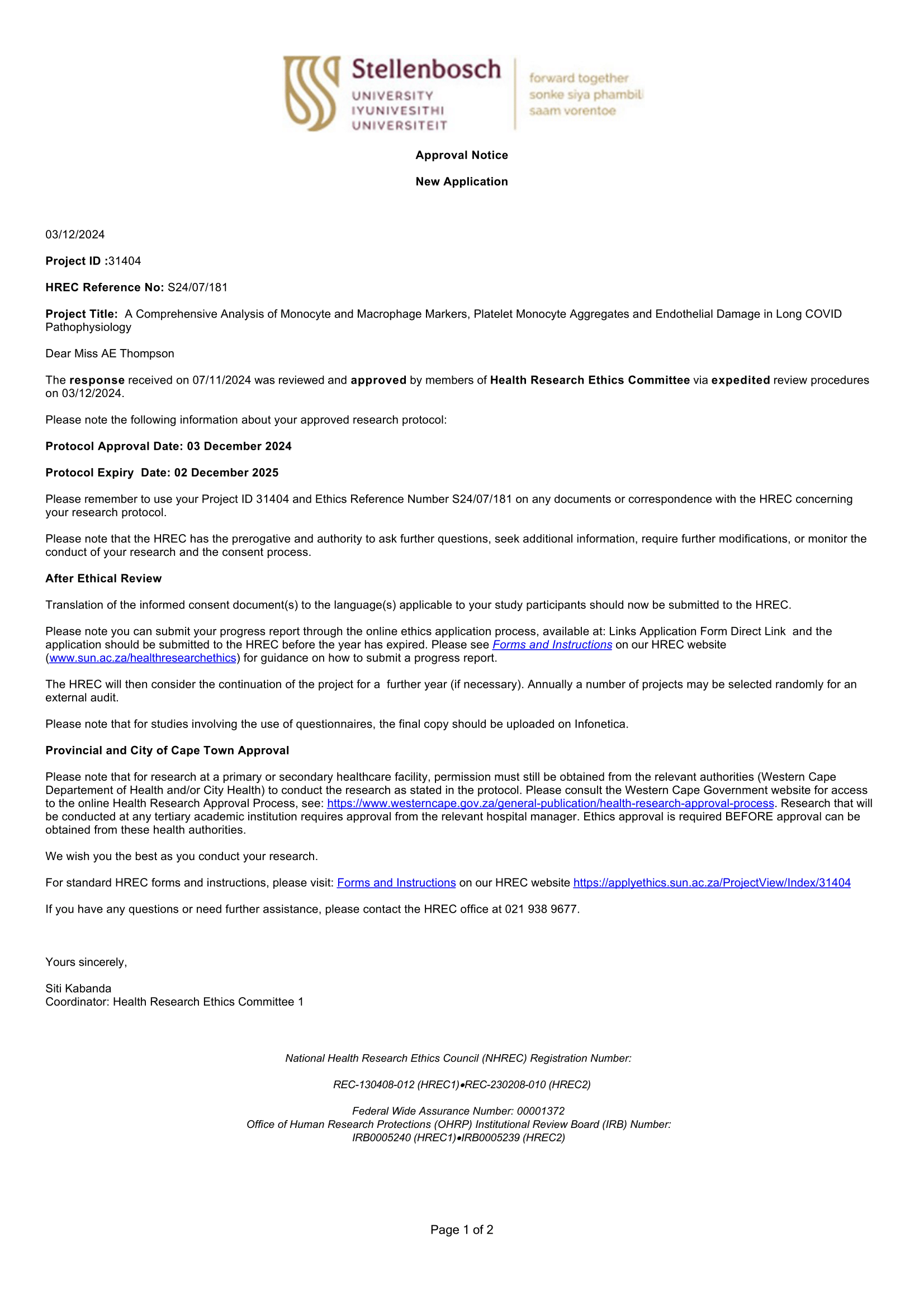

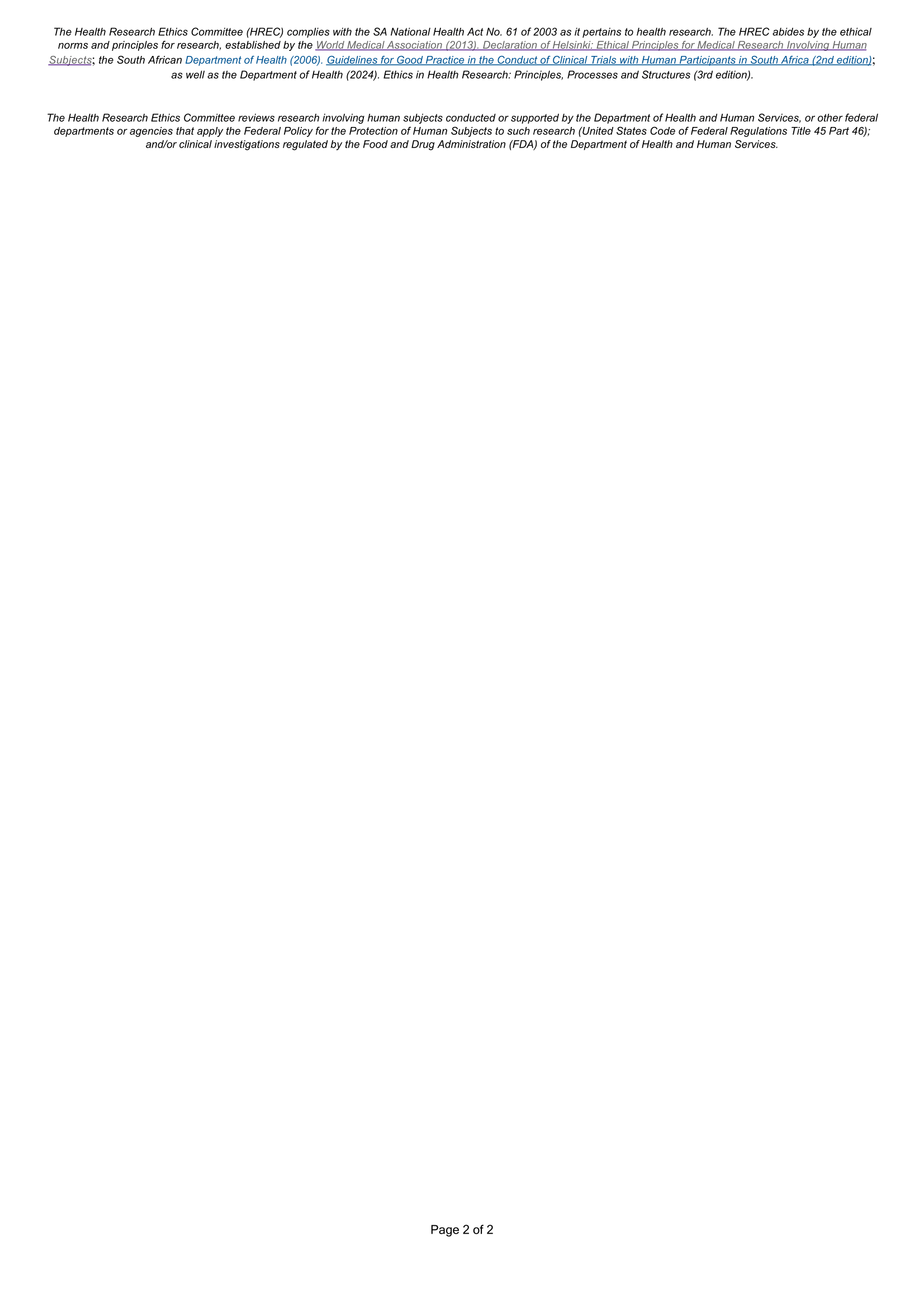
